# ITAF_45_ is a Pervasive *Trans* Acting Factor for Picornavirus Type II IRES Elements

**DOI:** 10.1101/2025.03.05.641721

**Authors:** Michael A. Bellucci, Mehdi Amiri, Stephen Berryman, Andia Moshari, Collins Oduor Owino, Rutger D. Luteijn, Tobias J. Tuthill, Yuri Svitkin, Graham J. Belsham, Frank JM van Kuppeveld, Nahum Sonenberg

**Author notes:** These authors contributed equally to this work (Michael A. Bellucci, Mehdi Amiri).

## Abstract

Viruses have evolved elaborate mechanisms to hijack the host mRNA translation machinery to direct viral protein synthesis. Picornaviruses, whose RNA genomes lack a cap structure, inhibit cap-dependent mRNA translation, and utilize an internal ribosome entry site (IRES) in the RNA 5′-UTR to recruit the 40S ribosomal subunit. IRES activity is stimulated by a set of host proteins termed IRES *trans*-acting factors (ITAFs). The cellular protein ITAF_45_ (also known as PA2G4 and EBP1) was identified as an essential ITAF for foot-and-mouth disease virus (FMDV), with no apparent role in cell-free systems for the closely related viruses harboring similar IRES elements such as encephalomyocarditis virus (EMCV) and Theiler’s murine encephalomyelitis virus (TMEV). Here, we demonstrate that ITAF_45_ is a pervasive host factor within cells for picornaviruses containing a Type II IRES. CRISPR/Cas9 knockout of ITAF_45_ in several human cell lines conferred resistance to infection with FMDV, EMCV, TMEV, and equine rhinitis A virus (ERAV). We show that ITAF_45_ enhances initiation of translation on type II IRESs in cell line models. This is mediated by the C-terminal lysine-rich region of ITAF_45_ known to enable binding to viral RNA. These findings challenge previous reports of a unique role for ITAF_45_ in FMDV infection, positioning ITAF_45_ as a promising antiviral target for various animal viruses and emerging human cardioviruses.

## INTRODUCTION

The canonical mechanism of mRNA translation initiation in eukaryotic cells entails the recruitment of the small ribosomal subunit (40S) to the mRNA via the 5′-terminal cap structure (m^7^GpppN, where m represents a methyl group, and N is any nucleotide), followed by ribosome scanning of the 5′-untranslated region (UTR) and recognition of an initiation codon^1^. Viruses do not encode translation components, therefore, they require the host translational machinery to sustain their life cycle^2^. Poliovirus was the first virus described to cause a shut-off of host protein synthesis^3^. The single-stranded positive-sense RNA of picornavirus genomes lack a cap structure^4,5^ and contain a long (400-1200 nt) 5′-UTR harboring a highly structured element termed an internal ribosome entry site (IRES)^6,7^. The IRES recruits the 43S initiation complex, which consists of the small 40S ribosomal subunit in association with eukaryotic initiation factors (eIFs)^8^. Picornavirus IRESs are classified into six types based on shared structural features, host factor requirements, and the initiation mechanism they employ^9,10^. Picornavirus Type I (*Enterovirus/Rhinovirus*) and Type II (*Cardiovirus/Aphthovirus*) groups differ most notably in that the latter recruits the 40S at, or close to, the initiation codon without the need for scanning^8^. Picornaviral IRES-mediated translation initiation is facilitated by cellular IRES *trans-*acting factors (ITAFs), which bind the IRES to engender conformational remodelling which promotes 43S binding^11,12^. ITAF requirements for different viruses are diverse, due in part to differential cell and tissue expression, which render ITAFs important determinants of host and tissue tropism^13^.

Proliferation-associated 2G4 (PA2G4), also referred to as ErbB3-binding protein 1 (EBP1) and IRES *trans*-acting factor 45 (referred throughout as ITAF_45_) is an evolutionarily conserved protein with multiple functions ranging from cell proliferation to embryonic development^14-17^. Two major isoforms exist, p48 and p42, wherein p42 lacks 54 amino acids at the N-terminus due to alternative splicing^17^. The p48 isoform is more abundantly expressed in most mammalian cells and is generally considered to have oncogenic properties in contrast to p42’s tumour-suppressive function^18^. p48 was first reported as an ITAF in a reconstituted cell-free translation system translation system in which 48S complex formation on the FMDV IRES required the mouse homologue of ITAF_45_ (murine proliferation-associated protein (*Mpp1*)^19^. While sequence similarity amongst the Type II IRES family can be about 50%, the distinct sequences largely retain a shared conserved secondary structure^20^. For translation initiation on IRES elements to occur, the recruitment of the 43S subunit, eIFs, and ITAFs is required to produce the 48S initiation complex^21^. Despite the similarities between the Type II IRES elements, ITAF_45_ was found to be dispensable for the assembly of 48S initiation complex formation on the EMCV and TMEV IRES in cell-free assays^19^. Subsequently, the requirement of FMDV IRES-driven translation for ITAF_45_ using various *in vitro* assays was corroborated^22-25^. siRNA-mediated depletion of ITAF_45_ in HEK-293 cells dramatically reduced FMDV IRES translation without affecting that of EMCV^25^. In sharp contrast, two recent independent CRISPR screens identified ITAF_45_ as a potential EMCV host factor, but in neither case was it validated^26,27^. Consequently, we sought to revisit the role of ITAF_45_ in RNA translation for EMCV and other Type II IRES-containing picornaviruses.

To address the discrepancy, we generated ITAF_45_ knockout (KO) cells using CRISPR-Cas9. We demonstrate that ITAF_45_ is a host factor for EMCV infection of human cells where it plays a critical role in stimulating IRES-driven translation. This requires the C-terminal lysine-rich region which has been shown to bind viral RNA^25^. Importantly, we demonstrate that ITAF_45_ is also required for infection of FMDV, TMEV, and ERAV, indicating that ITAF_45_ serves as a common host factor for picornaviruses containing a Type II IRES. This work positions ITAF_45_ as an important host factor that could serve as a therapeutic target for emerging human Cardiovirus diseases.

## RESULTS

Two recently published CRISPR screens have identified ITAF_45_ to be a potential EMCV host dependency factor^26,27^. To substantiate the claim for a role of ITAF_45_ in EMCV infection, we generated ITAF_45_ knockout (KO) H1-HeLa cells (Fig S1A, Fig. 1A), which were infected with viruses harboring different types of IRES elements (or lacking an IRES entirely) (Fig. 1B). These included species of the *Enterovirus* genus, EMCV (*Cardiovirus)*, and the unrelated vesicular stomatitis virus (*Rhabdovirus*), with a negative sense RNA genome (Fig. 1B). Cells were infected at high multiplicity of infection (MOI) and monitored until complete cytopathic effect (CPE) was observed in the control cells. As expected, ITAF_45_ depletion had no effect on enterovirus or rhabdovirus infection as shown by crystal violet staining of viable cells (Fig. 1B). Remarkably, depletion of ITAF_45_ conferred dramatic resistance to EMCV infection (Fig. 1B). This phenotype was specific to ITAF_45_ as restoring expression of full-length ITAF_45_ (with a C-terminal MYC-DDK tag) in the KO cells re-sensitized them to infection (Fig. 1B). Furthermore, infection of ITAF_45_^KO^ cells with EMCV showed a ∼300-fold reduction in viral RNA production following one cycle of infection (Fig. 2A). Virus yield assays using the supernatant of EMCV-infected control and ITAF_45_ ^KO^ Λ a ∼10,000-fold reduction in the titer of infectious virus that was rescued by restored expression of ITAF_45_ in the KO cells (Fig. 2B). To determine whether the phenotype was reproduced in other cell lines, we generated ITAF_45_ ^KO^ eHAP1 (haploid human cell line derived from the chronic myelogenous leukemia (CML) cell line KBM-7 (Fig. S2A). Upon infection of eHAP1 ITAF_45_ ^KO^ cells at high MOI, the cells were resistant to EMCV replication which was rescued upon restoration of ITAF_45_ expression using ITAF_45_-MYC-DDK (Fig. S2B). Therefore, ITAF_45_ is a required host factor for EMCV infection in different human cell lines.

**Figure 1.**
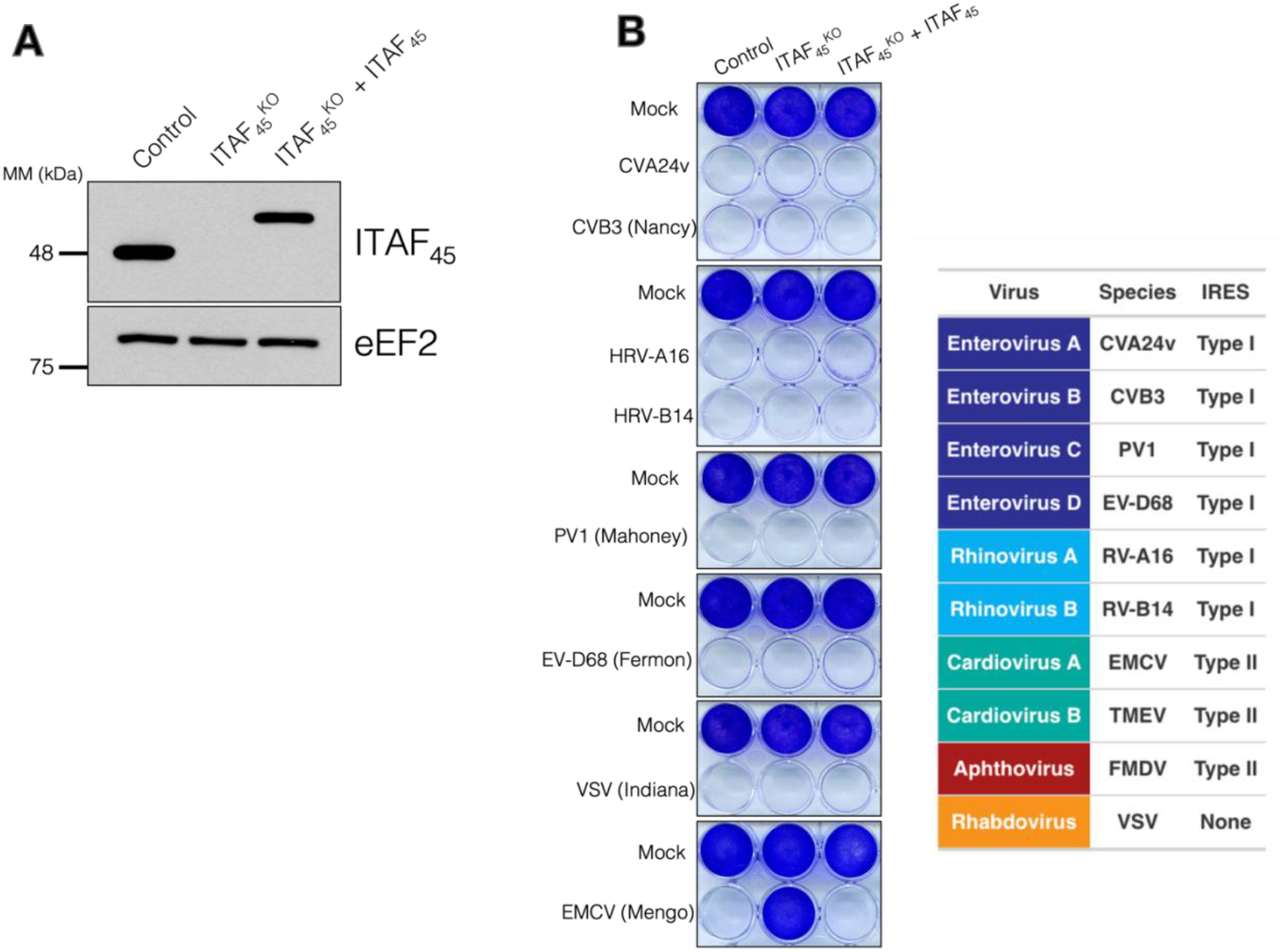
ITAF_45_ is a key host factor for EMCV infection. **A)** Immunoblot of control, ITAF_45_^KO^, and ITAF_45_^KO^ + MYC-DDK-tagged ITAF_45_ H1-HeLa cells. **B)** Crystal violet staining of viable cells infected with indicated virus with “mock” representing uninfected cells. (**Also B**): Categorization and taxonomy of viruses used and the type of IRES they contain in their 5′-UTR.

**Figure 2.**
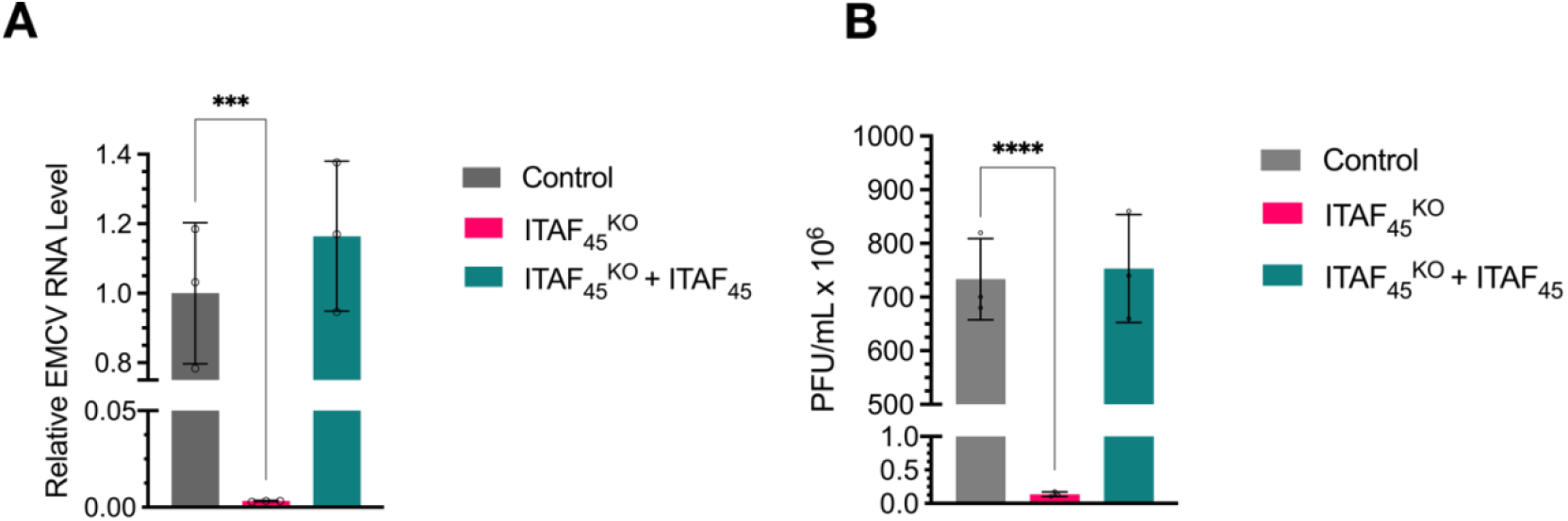
Depletion of ITAF_45_ restricts EMCV infection. **A)** RT-qPCR quantification of EMCV RNA in control, ITAF_45_^KO^, and ITAF_45_^KO^ + ITAF_45_ infected with EMCV (*Mengo* strain) at a multiplicity of infection (MOI) of 10 for 7 hours. **B)**. Plaque assay quantification of infectious EMCV produced from control, ITAF_45_^KO^, and ITAF_45_^KO^ + ITAF_45_ infected with EMCV (*Mengo* strain) at an MOI of 10 for 24 hours. Datasets represent means ± s.d. (n = 3 independent biological replicates). All P values were determined by ordinary one-way ANOVA using GraphPad Prism (GraphPad Software) with Dunnet correction. ***P < 0.0002 and ****P < 0.0001.

Next, we sought to investigate the possibility that ITAF_45_ was involved in viral entry, independent of IRES-mediated translation. To this end, we transfected EMCV RNA transcripts containing a *Gaussia* luciferase reporter (EMCV-GLuc) downstream of the EMCV RNA IRES into control and ITAF_45_^KO^ cells using liposomes, thus bypassing the virus entry mechanism. At 7 h post-transfection, we observed a ∼20-fold reduction in *Gaussia* luciferase activity in the ITAF_45_^KO^ as compared to control cells (Fig. 3A), establishing that ITAF_45_ is acting downstream of viral entry. We next investigated whether ITAF_45_ was required for viral translation by first transfecting control or ITAF_45_^KO^ cells with a capped and polyadenylated bicistronic reporter mRNA containing the EMCV IRES between firefly and *Renilla* luciferase cistrons (Fig. 3B). Cells were lysed 6 h post-transfection when we observed a 4-fold reduction in IRES-mediated *Renilla* luciferase translation in ITAF_45_^KO^ cells, but no change in cap-dependent translation (Fig. 3B). To bolster these results, we next performed a synchronized infection assay on control and ITAF_45_^KO^ cells. Virus was first rescued in BHK-21 cells from the EMCV-GLuc RNA transcript mentioned above. Using cycloheximide (CHX), a potent translation inhibitor, and EMCV-GLuc virus, this assay was employed to elucidate whether depletion of ITAF_45_ affects the viral translation stage (<3 hr post-infection) or at later stages such as viral replication (Fig. 3C). The exponential increase in luciferase activity observed in infected control cells was abolished by CHX (Fig. 3C). Strikingly, the luciferase signal observed in ITAF_45_^KO^ cells without CHX was equal to control cells treated with CHX (Fig. 3C). These experiments demonstrate that ITAF_45_ enhances EMCV IRES-mediated translation in human cell lines and is required to support virus infection.

**Figure 3.**
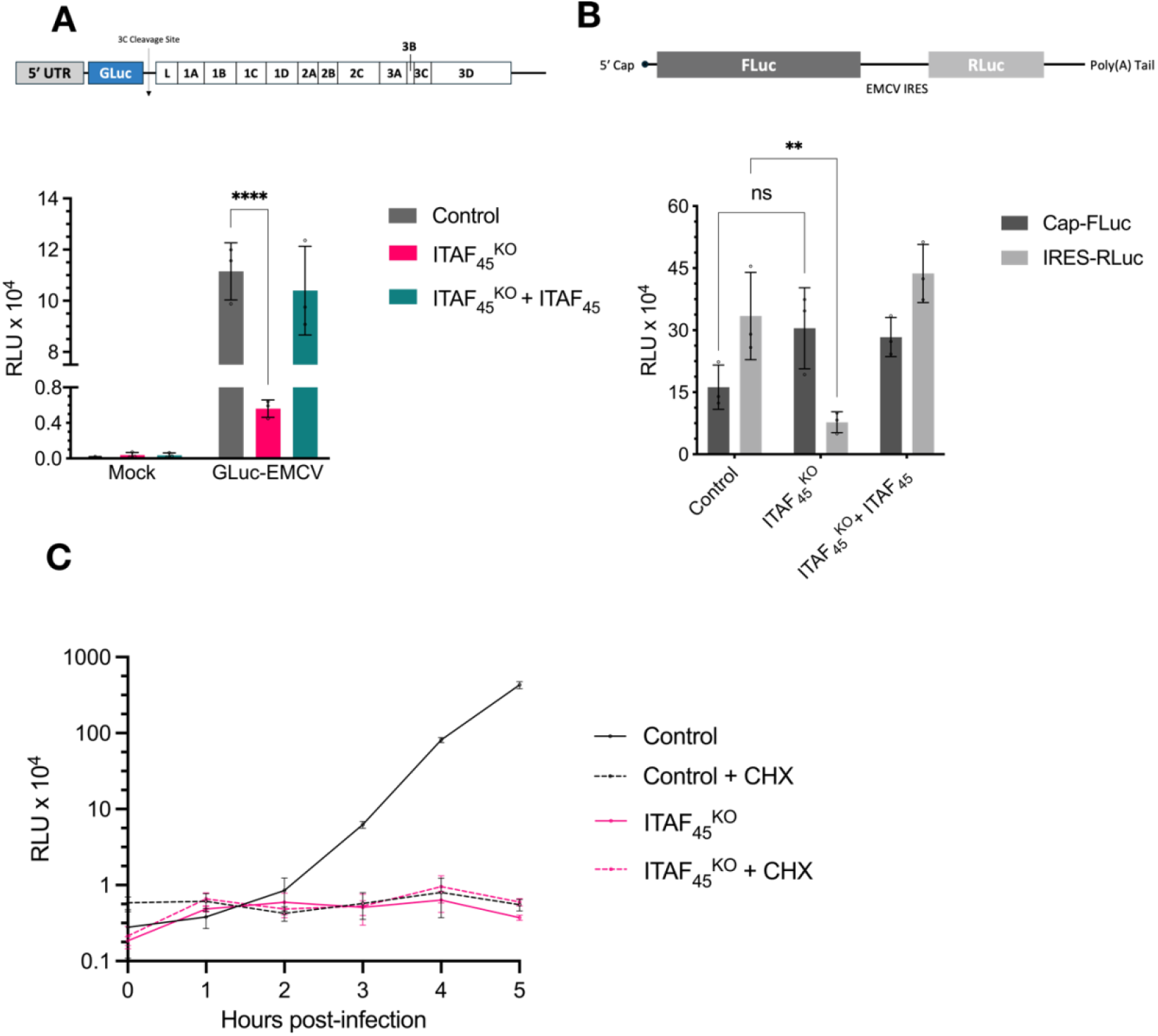
ITAF_45_ stimulates the translation of EMCV RNA in human cell lines. Top: Schematic of the full-length EMCV (*Mengo* strain) genome containing *Gaussia* luciferase downstream of the 5′-UTR followed by a 3C^pro^ cleavage site (EMCV-GLuc). Bottom: EMCV-GLuc expression levels in control, ITAF_45_^KO^, and ITAF_45_^KO^ + ITAF_45_ cells transfected with 1 μg of EMCV-GLuc RNA. **B**. Top: Schematic of the capped and polyadenylated EMCV IRES bicistronic reporter mRNA used in which the EMCV IRES is between a *Firefly* luciferase cistron (cap-dependent translation) and a *Renilla* luciferase cistron (IRES-mediated translation). Bottom: *Firefly* and *Renilla* luciferase expression levels in relative light units (RLU) in control, ITAF_45_^KO^, and ITAF_45_^KO^ + ITAF_45_ cells transfected with 400 ng of the EMCV bicistronic reporter mRNA. Cells were harvested at 6 hr post-transfection. Background signal from non-transfected cells were subtracted from the measurements. **C**. Control and ITAF_45_^KO^ cells infected with EMCV-GLuc virus at an MOI of 20 in the presence of 0.1% DMSO or 125 µM cycloheximide (CHX). Datasets represent means ± s.d (n = 3 independent biological). All P values were determined by two-way ANOVA using GraphPad Prism (GraphPad Software) with Dunnet correction. ***P < 0.0002 and ****P < 0.0001.

These data show that ITAF_45_ acts as a critical ITAF for EMCV, in addition to what was previously reported for its role in FMDV infection. To date, the role of ITAF_45_ for FMDV RNA translation has only been studied either in cell-free translation reconstitution assays or using transfection of artificial constructs bearing the FMDV IRES. Thus, we sought to confirm that ITAF_45_ is also required for viral translation in FMDV-infected cells. eHAP1 control and ITAF_45_ ^KO^ cells were infected with FMDV (*O1 Kaufbeuren* strain) at high MOI and imaged every hour to assess CPE development (Fig. 4A). At 12 - 15 h post-infection, control cell confluency was significantly reduced with most cells displaying marked CPE, while ITAF _45_^KO^ cells remained unaffected, clearly establishing that full-length FMDV requires ITAF_45_ during infection in a cell line model (Fig. 4A). Furthermore, when transfected with a GFP-containing FMDV replicon in which the capsid coding region was replaced with a GFP reporter, the GFP signal observed in control cells was lost in ITAF_45_ ^KO^ cells and did not surpass the levels observed in infected control cells that were co-treated with guanidine hydrochloride (viral replication inhibitor) (Fig. 4B). In addition to FMDV, we explored the role of ITAF_45_ in TMEV (*GDVII* strain) replication and ERAV titration (Fig. 4C and 4D). As with FMDV and EMCV, ITAF_45_^KO^ cells failed to support TMEV infection in human cell lines as measured by RT-qPCR, while a 4-log reduction in virus yield was observed during an ERAV infection. These results demonstrate a broad role for ITAF_45_ for type II IRES-containing viruses of the cardio- and aphtho-virus families.

**Figure 4.**
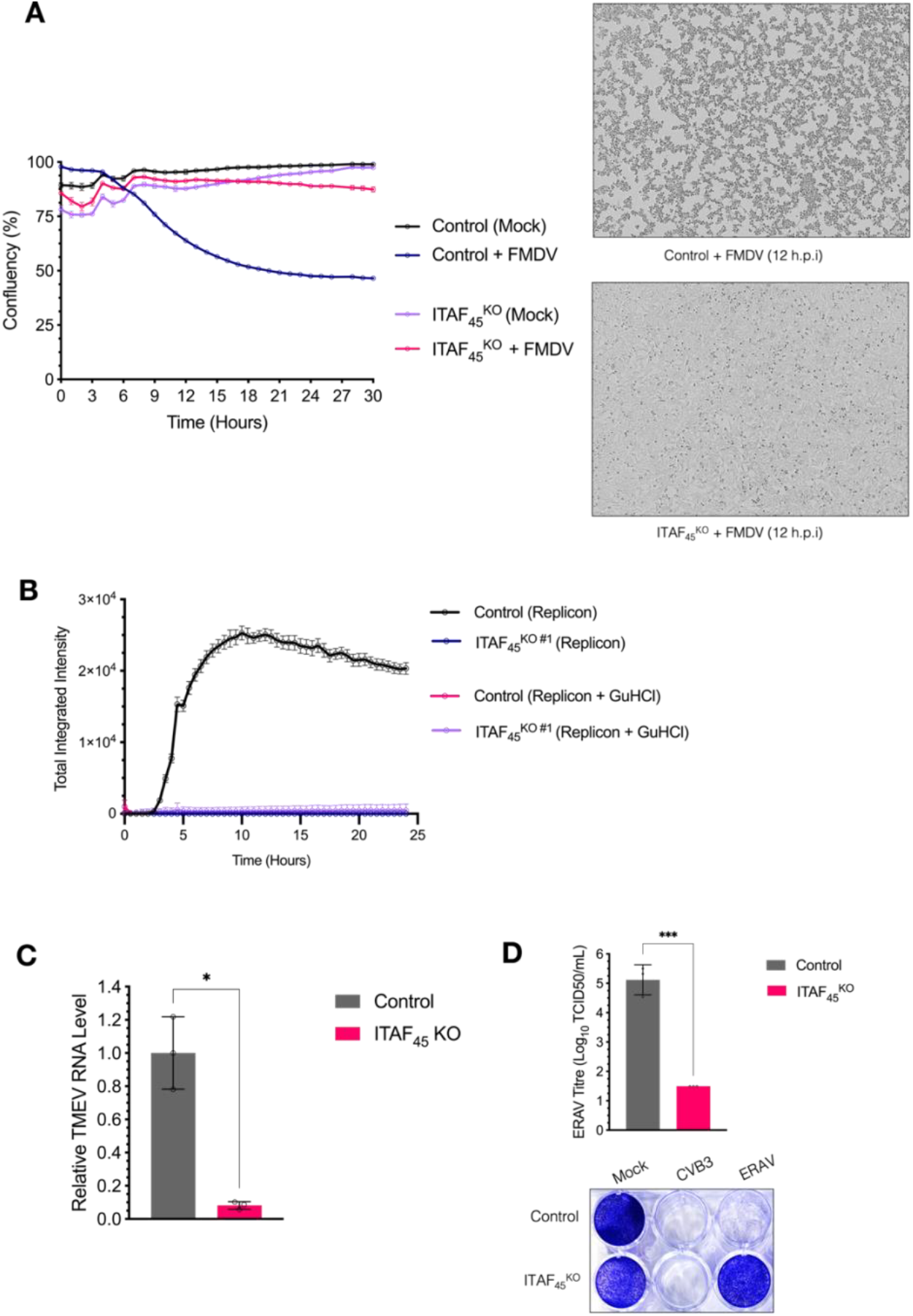
ITAF^45^ is required for viruses containing a type II IRES. **A**. Left: eHAP1 control and ITAF_45_^KO^ cells were infected with FMDV (*O1 Kaufbeuren*) and cultured in an incucyte to monitor the rate of cytopathic effect (CPE) development via measurement of cell confluency. The average confluency of three biological replicates was plotted against time using GraphPad Prism (GraphPad Software). Right: Representative image of cell monolayers at 12 hours post-infection. **B.** Replication of FMDV in eHAP1 control and ITAF_45_^KO^ cells was assessed via transfection of 90 ng of an FMDV subgenomic replicon wherein the capsid protein coding regions were replaced with a GFP reporter. Replication was assessed using the Incucyte S3 2019B ver2 software to quantify the total integrated green intensity per well using a cutoff value of 0.5. **C**. eHAP1 control and ITAF_45_^KO^ cells were infected with TMEV (*GDVII* strain) at an MOI of 50 and lysed at 24 hrs post-infection to quantify viral RNA using RT-qPCR. **D. Top:** eHAP1 control and ITAF_45_^KO^ cells were used to titrate ERAV stock where the end-point titres were calculated using the Spearman Kärber method. **Bottom:** Crystal violet staining of viable cells inoculated with indicated virus with “mock” representing untreated cells. TMEV and ERAV dataset represent means ± s.e.m. (n = 3 independent biological replicates). P values were determined by a paired T-test using GraphPad Prism (GraphPad Software) *P < 0.0332.

The C-terminal lysine-rich region of ITAF_45_ is required for its nucleic acid-binding properties, including to the EMCV and FMDV IRESs^25^. To investigate whether the C-terminal lysine-rich region is required for IRES-driven translation in cells, we constructed ITAF_45_ ^KO^ cell lines that stably express a MYC-DDK FLAG-tagged truncated ITAF_45_ (amino acids 1-360) lacking the lysine-rich region or a full-length ITAF_45_ containing a modified lysine-rich region (^365^KKKKKKKSKT^376^ changed to ^365^AAAAAAASAT^376^) (Fig. 5A). Expression of either mutant protein, in contrast to the full-length WT ITAF_45_, in ITAF_45_ ^KO^ cells failed to rescue virus replication, indicating that the C-terminal lysine-rich region of ITAF_45_ is required for EMCV infection (Fig. 5B).

**Figure 5.**
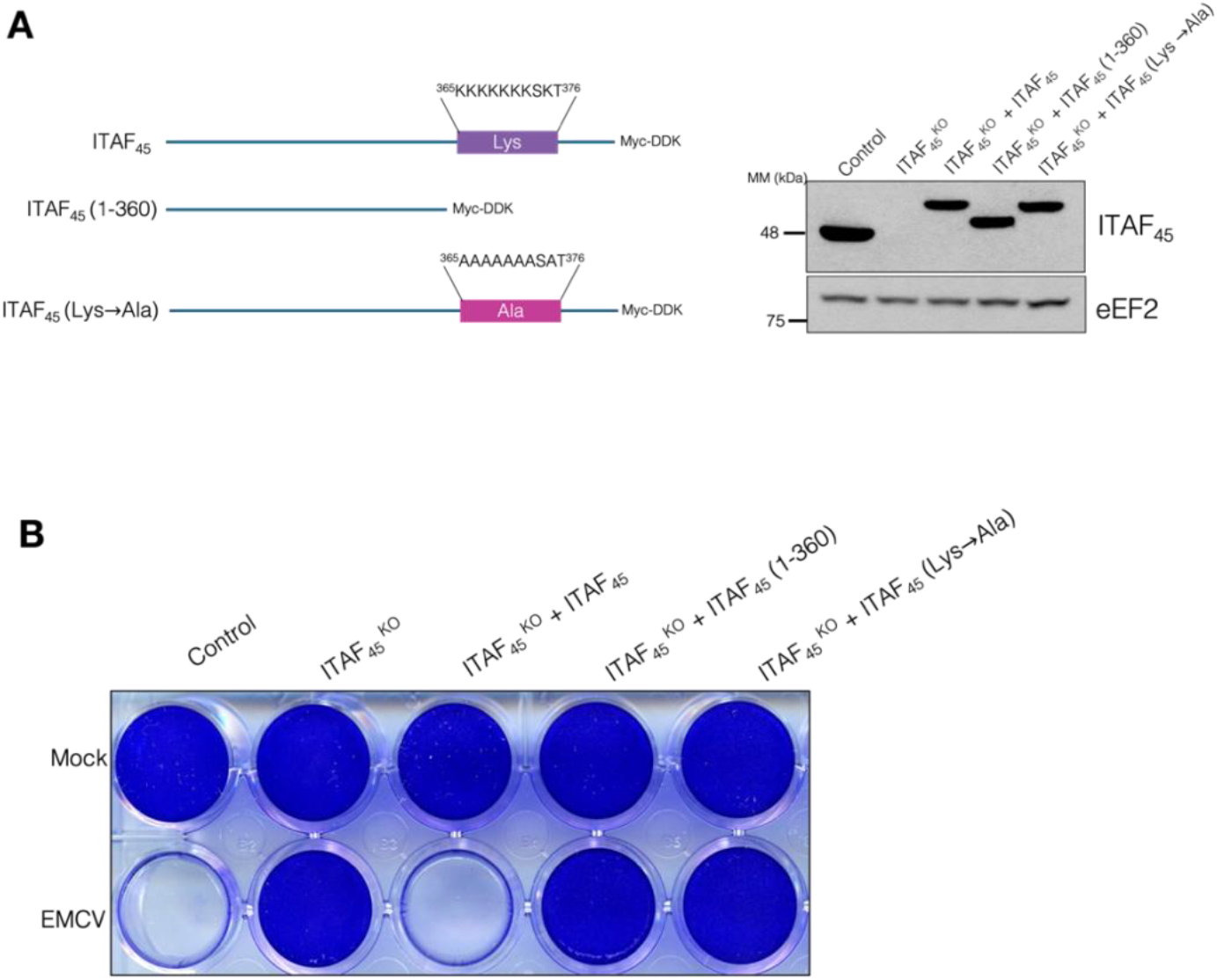
C-terminal lysine-rich region of ITAF_45_ is required for efficient EMCV infection. **A. Left:** Schematic of ITAF_45_ constructs stably expressed in H1-HeLa ITAF_45_^KO^ cells containing C-terminal MYC-DDK tags. **Right:** Immunoblot of H1-HeLa control, ITAF_45_^KO^, and ITAF_45_^KO^ + MYC-DDK-tagged construct cell lines. **B**. Crystal violet staining of viable cells infected with EMCV (*Mengo* strain) at an MOI of 10 for 24 hrs with “mock” representing uninfected cells.

## DISCUSSION

Here, we demonstrate that ITAF_45_ is as an important host factor for type II IRES-containing picornaviruses. We show that upon CRISPR/Cas9-mediated loss of ITAF_45_ expression in two human cell lines, infection with EMCV is severely restricted. Restored expression of the full-length p48 isoform of ITAF_45_ reverted this phenotype, while expression of modified ITAF_45_ with a disrupted C-terminal lysine-rich region did not. This is consistent with previous data that identified the C-terminal lysine-rich region as the essential motif which imparts the nucleic acid binding activity of ITAF_45_.^25^. The p42 isoform was weakly expressed in the parental H1-HeLa and eHAP1 cells, consistent with reports that the p42 isoform is not detectable in cancer cell lines due to its ubiquitin-mediated degradation via BRE1^28,29^.

Using an *in vitro* bicistronic reporter assay and a synchronized infection assay in cells, we demonstrated that ITAF_45_ greatly stimulates the IRES-mediated translation of the EMCV genome. Furthermore, using full-length FMDV, TMEV, and ERAV, we showed that ITAF_45_ is also necessary for efficient replication of each virus in cell line models. These results are in stark contrast to the earlier conclusion that ITAF_45_ functions exclusively in FMDV RNA translation. In previous reports, footprinting and toeprinting assays of the FMDV IRES showed that ITAF_45_ bound the base of domain I of the FMDV IRES at a region with very high similarity to the corresponding region of the EMCV IRES^19,23,30,31^. In addition, while ITAF_45_ was shown to crosslink to the EMCV and TMEV IRES following UV irradiation^19^, and could bind the EMCV IRES *in vitro*^25^, initiation on either IRES was independent of ITAF_45_. A potential explanation for this discrepancy is the different *in vitro* constructs used. For example, it was reported that within an *in vitro* context, one extra A nucleotide in the EMCV IRES A-rich bulge conferred translational dependence on PTB (polypyrimidine tract binding protein, another well-characterized ITAF) which is otherwise not required by the WT IRES^32^. This, however, was shown to be highly variable depending on the sequence of the IRES itself and the type of reporter downstream of the IRES (viral or heterologous cistron)^32^. In our studies, we utilized both a reporter mRNA with the wild-type EMCV IRES driving translation of non-viral cistrons as described by Palmenberg^33^, and corroborated those results using infectious virus containing the WT IRES sequence in cell line models.

The IRES was first discovered in poliovirus and EMCV RNAs^6,7^. EMCV and FMDV pose a major risk to agricultural and livestock industries by infecting a diverse array of species^34-37^. Very few cases of symptomatic EMCV infections in humans have been reported, although serological studies have shown that particularly in developing countries and amongst populations exposed to livestock, EMCV is a widely circulating virus^38-41^. Importantly, the emergence of novel human *Cardioviruses* such as Saffold virus has elicited great interest in understanding the life cycles of these viruses to prepare for possible future outbreaks^42-46^.

Importantly, we showed using a bicistronic reporter mRNA containing the EMCV IRES between heterologous reporters (luciferase cistrons)^33^ that IRES-dependent translation was impaired 4-fold in ITAF_45_ ^KO^ cells. It is noteworthy that the only prior cell-based assay investigating ITAF_45_’s role in EMCV infection (using an EMCV IRES construct transfected into HEK-293 cells with siRNA-mediated ITAF_45_ depletion) found no impact on EMCV translation^25^. While there was no difference in the IRES-dependent cistron used between our experiments, the reported RNA transfection was performed for 24 hrs (using a non-replicating mRNA) in contrast to our experiment with an endpoint of 6 hrs post-transfection. In addition, since siRNA confers a transient and partial knockdown of protein levels, it is conceivable that this was not sufficient to confer the phenotype we observe using a CRISPR/Cas9 complete knockout. This work further highlights the critical importance of the control of RNA translation in the viral life cycle. As shown with the IRES reporter assays, relatively modest repression of EMCV IRES translation in ITAF_45_ ^KO^ cells causes a drastic reduction in viral replication, presumably due to the importance of multiple cycles of translation that occur immediately following infection.

We showed the requirement of ITAF_45_ for FMDV, EMCV, TMEV, and ERAV infection in cell lines. This corroborated previous *in vitro* experiments using artificial FMDV reporters and importantly, dispelled the previously mischaracterized absence of a role for ITAF_45_ in EMCV and TMEV RNA translation. With the recent isolation of emerging human cardioviruses that exhibit a high degree of sequence similarity with the 5′-UTR of TMEV RNA^47^, this work documents a possible therapeutic target to combat picornaviruses harbouring type II IRES elements.

## METHODS

### Cells and Reagents

HEK293T, eHAP1, BHK-21, and H1-HeLa cells were obtained from the American Type Culture Collection (ATCC). HEK293T, BHK-21, and H1-HeLa cells were maintained in Dulbecco’s modified Eagle’s medium supplemented with 10% fetal bovine serum and 1% penicillin/streptomycin. eHAP1 cells were maintained in Iscove’s modified Dulbecco’s medium supplemented with 10% fetal bovine serum and 1% penicillin/streptomycin. For FMDV experiments, eHAP1 cells were maintained in Virus Growth Media with reduced (1%) serum.

### Viruses and Infectious Clones

Infectious clones of encephalomyocarditis virus (EMCV *Mengo* strain), coxsackievirus B3 (CVB3 *Nancy* strain), CVA24v, enterovirus D68 (EV-D68 *Fermon* strain), EMCV-GLuc, human rhinovirus (HRV)-A16 and HRV-B14 were obtained from Frank Kuppeveld and poliovirus 1 (PV1 *Mahoney* strain) from Eckard Wimmer. Vesicular stomatitis virus (VSV *Indiana* strain) was obtained from ATCC (VR-1421). EMCV, EMCV-GLuc, and foot-and-mouth disease virus (FMDV *O1 Kaufbeuren* strain) infectious clones were rescued and titrated on BHK-21 cells. Equine rhinitis A virus (ERAV *NM11/67*) was obtained from David Rowlands and Toby Tuthill (University of Leeds, Leeds, United Kingdom). CVB3 and PV1 infectious clones were rescued and titrated on H1-HeLa cells. EV-D68 infectious clone was rescued and titrated on RD cells. HRV infectious clones were rescued and titrated on H1-HeLa cells. Infectious clones (EMCV and EMCV-GLuc) were rescued as viruses by first linearizing the plasmids with *BamHI* restriction enzyme (Thermo Fisher Scientific). CVB3, CVA24v, and EV-D68 were linearized with *SalI* (Thermo Fisher Scientific), PV1 and HRV-B14 were linearized with *MluI* (Thermo Fisher Scientific), and HRV-A16 was linearized with *SacI* (Thermo Fisher Scientific). The linearized plasmids were *in vitro* transcribed using HiScribe® T7 High Yield RNA Synthesis Kit (New England Biolabs) and purified using an RNA Clean and Concentrator Kit (Zymo Research). 5 μg of derived RNA was transfected into the cell lines mentioned above using Lipofectamine-2000 (Thermo Fischer Scientific). Between 24-72 hours post-transfection (when cytopathic effect was >90%), rescued virus was harvested from the supernatant and cell lysates following three freeze-thaw cycles and high-speed centrifugation. The resulting passage 0 virus was added to the appropriate cell lines and harvested in the same manner followed by titration on the appropriate cell lines using plaque assays as described by Rueckert ^48^. Passage 1 viral stocks were used for all experiments unless otherwise stated.

### Generation of Cell Lines

To generate ITAF_45_ knockout cell lines (ITAF_45_^KO^), H1-HeLa and eHAP1 cells expressing spCas9 were generated. HEK293T cells were first transfected with Lenti-Cas9-2A-Blast (Addgene #73310) along with psPAX2 (Addgene #12260), pMD2.G (Addgene #12259), and Lipofectamine-2000 (Thermo Fischer Scientific) to generate lentivirus. H1-HeLa and eHAP1 cells were then transduced with Cas9-2A-Blast lentivirus followed by blasticidin selection. The selected cells were single cell sorted using flow cytometry to obtain clonal populations with sufficient Cas9 expression. Then, two independent guide RNAs targeting different exons of ITAF_45_ (ACAGGAGCAAACTATCGCTG and GGGTTGGCACCTACTTCTGC) or non-targeting sequences (ACGGAGGCTAAGCGTCGCAA and CGCTTCCGCGGCCCGTTCAA) were cloned into a lentiviral expression plasmid as described in Addgene #154194. The resulting plasmid was then transfected into HEK293T cells to generate lentivirus. Clonal populations of H1-HeLa-Cas9 and eHAP1-Cas9 cells were transduced with the control or ITAF_45_ ^KO^ lentivirus, selected with puromycin, and sorted using flow cytometry to obtain clonal populations. Knockouts were validated using Western blotting and/or Sanger sequencing. To restore expression of ITAF_45_ in the ITAF_45_ ^KO^ cell lines, synthetic oligos corresponding to the cDNA of the C-terminal MYC-DDK-tagged ITAF_45_ or the C-terminal mutant constructs described above were obtained from Integrated DNA Technologies and cloned into a lentiviral construct (Addgene #104995). The lentiviral transfection and transduction procedures were carried out in the same manner as previously mentioned.

### Virus Screening and Crystal Violet Staining

Control, ITAF_45_^KO^, and ITAF_45_ ^KO^ + ITAF_45_ cells were seeded onto 12-well plates. The following day, cells were incubated with CVA24v, CVB3, RV-B14, RV-A16, PV-1, EV-D68, VSV, and EMCV at high MOI in culture medium for 30 min at 37ºC (CVB3, PV-1, VSV, and EMCV) or for 1 hr at 33ºC (CVA24v, RV-B14, RV-A16, and EV-D68). Following virus adsorption, cells were washed three times with PBS and replenished with fresh media. Cells were incubated for 24 – 48 h and then fixed and stained using 0.5% crystal violet in methanol.

### FMDV Experiments

For FMDV infections to assess CPE development, eHAP1 control and ITAF_45_^KO^ cells were seeded in 96-well plates. Wells were washed once with Virus Growth Media (VGM: normal cell culture medium with reduced (1%) serum), and infected with FMDV (*O1 Kaufbeuren*) in a volume of 50 μL. After 1 hr at 37°C, the virus inoculum was replaced with 150 μL of VGM. The infection was continued at 37°C and the cells in each well were imaged at 1 hr intervals (10 x objective, 4 images per well) for 30 hrs using the Incucyte S3 live cell analysis system (Essen Biosciences). Cell confluency was calculated from the images using the Incucyte S3 2019B ver2 software with default software settings. To assess the rate of CPE development, the average confluency of three replicate wells was plotted against time. Replication of FMDV in eHAP1 cells was assessed using a previously described subgenomic replicon in which the capsid protein coding region was replaced with a fluorescent reporter sequence (ptilosarcus GFP; ptGFP). The replicon plasmid was linearized using *AscI* (New England Biolabs) and purified using the GFX Illustra DNA and Gel Band Purification Kit (Cytiva). 1 μg of linearized DNA was transcribed *in vitro* using a Megascript T7 Kit (Thermo) as per the manufacturer’s instructions, and RNA transcripts were purified from the reaction using the Megaclear Transcription Clean Up Kit (Thermo). eHAP1 control and ITAF_45_^KO^ cells were grown to 90% confluency in 96-well tissue culture plates and washed once with VGM followed by transfection with 90ng of replicon RNA per well using a TransIT-mRNA Transfection Kit (Mirus). VGM alone or VGM containing 3 mM guanidine hydrochloride (GuHCl) was then added along with the transfection mixture to each well. The plates were incubated at 37°C and the wells imaged every 0.5 h using an Incucyte S3 live cell analysis system to assess fluorescent protein expression (10x objective, 4 images per well). The images were analysed using the associated Incucyte S3 2019B ver2 software to quantify the total integrated green intensity per well using a cutoff value of 0.5. The total integrated intensity is the sum of all green fluorescence intensity values within the green objects in the image, multiplied by the pixel area. To assess replication, the average total integrated intensity of four replicate wells was plotted against time.

### Reverse-Transcription Quantitative PCR (RT-qPCR) for Quantification of Viral RNA

Control, ITAF_45_^KO^, and ITAF_45_ ^KO^ + ITAF_45_ cells were seeded onto 6-well plates in triplicate. The following day, cells were infected with EMCV at an MOI of 10 in culture medium for 30 min at 37ºC. Following the 30-minute virus adsorption, cells were washed three times with PBS and replenished with fresh media. At 7 h post-infection, cells were washed with PBS and harvested in TRIzol (Invitrogen). RNA was extracted using Direct-zol RNA miniprep kit (Zymo Research) according to the manufacturer’s protocol. Reverse transcription, with 1 μg of RNA was performed using the SuperScript III Reverse-Transcriptase Kit (Invitrogen, Cat#18080044) and Oligo(dT)_12-18_ (Invitrogen, Cat#18418012) according to the manufacturer’s protocol. Quantitative PCR was performed using SYBR Green (Invitrogen, Cat#4309155) and CFX Connect Real-Time PCR Detection System (Bio-Rad) equipment. The following primers were used (5′ to 3′ orientation): *EMCV For*: CCACAGAGGATTGGAAGCCA; *EMCV Rev*: GCACGCAAAACTGCCTGATA, *GAPDH For*: TGGGTGTGAACCATGAGAAG; *GAPDH Rev:* ATGGACTGTGGTCATGAGTC; *TMEV For*: AGCCCATCCACGATGAGCTT; *TMEV Rev*: CTGAAAAACCGACTGCACAGG

### EMCV IRES Bicistronic Reporter Assay

The EMCV IRES bicistronic reporter plasmid was a gift from Ann Palmenberg ^33^. The plasmid was first linearized with *BamHI* (Thermo Fischer Scientific) and *in vitro* transcribed using HiScribe® T7 High Yield RNA Synthesis Kit (New England Biolabs) with Anti-Reverse Cap Analog (ARCA) (New England Biolabs #S1411) followed by purification using the RNA Clean and Concentrator Kit (Zymo Research). The purified capped mRNA product was polyadenylated using *E. coli* Poly(A) Polymerase (New England Biolabs # M0276) followed by purification using the RNA Clean and Concentrator Kit (Zymo Research). Control, ITAF_45_ ^KO^, and ITAF_45_ ^KO^ + ITAF_45_ cells were seeded into 24-well plates in triplicate. The following day, cells were transfected with 400 ng of capped and polyadenylated reporter mRNA using Liptofectamine-2000 (Thermo Fischer Scientific). At 6 h post-transfection, cells were harvested using the Dual-Glo® Luciferase Assay System (Promega) according to the manufacturer’s protocol. The *Renilla* (IRES) and Firefly (Cap) luciferase activity levels were measured using the dual-glo setting of the Glomax 20/20 Luminometer (Promega). Measurements were normalized to background levels from non-transfected cells.

### Viral RNA Transfection

Control and ITAF_45_^KO^ cells were seeded into 24-well plates in triplicate. The following day, 1μg of *in vitro* transcribed EMCV-GLuc RNA was transfected into cells using Lipofectamine-2000 (Thermo Fischer Scientific). At 7 h post-transfection, cells were harvested using the *Renilla* Luciferase Assay system (Promega) according to the manufacturer’s protocol and measured using the luc-0-inj setting of the Glomax 20/20 Luminometer (Promega). Non-transfected control and ITAF_45_ ^KO^ cells were used as controls.

### Synchronized Infection Assay with EMCV-GLuc

Control and ITAF_45_ ^KO^ cells were seeded into 24-well plates in triplicate. The following day, the cells were first chilled on ice for 30 min, after which the cells were incubated with 125 μM cycloheximide (BioShop) or 0.1% DMSO for 1 hr on ice with or without EMCV-GLuc (MOI = 20). After 1 hr incubation to allow the virus to attach to the cells, the plates were incubated for 10 min at 37ºC to permit virus entry. The cells were then washed three times with PBS and replenished with media containing the aforementioned compounds. At the indicated time points, cells were harvested using the *Renilla* Luciferase Assay system (Promega) according to the manufacturer’s protocol and measured using the luc-0-inj setting of the Glomax 20/20 Luminometer (Promega).

### SDS-PAGE and Western Blotting

Unless otherwise stated, cells were lysed using RIPA buffer (50 mM Tris-HCl pH 7.4; 1% NP-40; 0.1% SDS; 150 mM NaCl; 2 mM EDTA) supplemented with protease and phosphatase inhibitors (Roche). Protein lysates were incubated at 4°C with constant rotation for 30 min followed by centrifugation at 14,000 rpm for 15 min at 4°C. The supernatants were collected for protein quantification using Bradford reagent (BioRad). Protein lysates were denatured in 5X loading buffer (250 mM Tris-HCl pH 6.8; 8% SDS; 0.2% w/v bromophenol blue; 40% v/v glycerol; 20% v/v β-mercaptoethanol), boiled, and subjected to 10% polyacrylamide gel electrophoresis followed by transfer onto a nitrocellulose membrane. Membranes were blocked with 5% skim milk solution in TBST. ITAF_45_ antibody was obtained from Novus Biologials and eEF2 antibody was obtained from Cell Signalling Technology.

## ACKNOWLEDGEMENTS

We thank Stephen Curry for his insightful advice and Ann Palmenberg for providing the EMCV bicistronic reporter vector. This work was funded by the Terry Fox Foundation/OHRIT, CIHR PJT-192041, and CIHR FDN-148423. The flow cytometry and cell sorting was performed in the Flow Cytometry Core Facility for flow cytometry and single cell analysis of the Life Science Complex and supported by funding from the Canadian Foundation for Innovation. FMDV experiments were performed within the high containment labs at The Pirbright Institute and ERAV experiments were performed by the Kuppeveld Lab.

---

**Figure S1.**
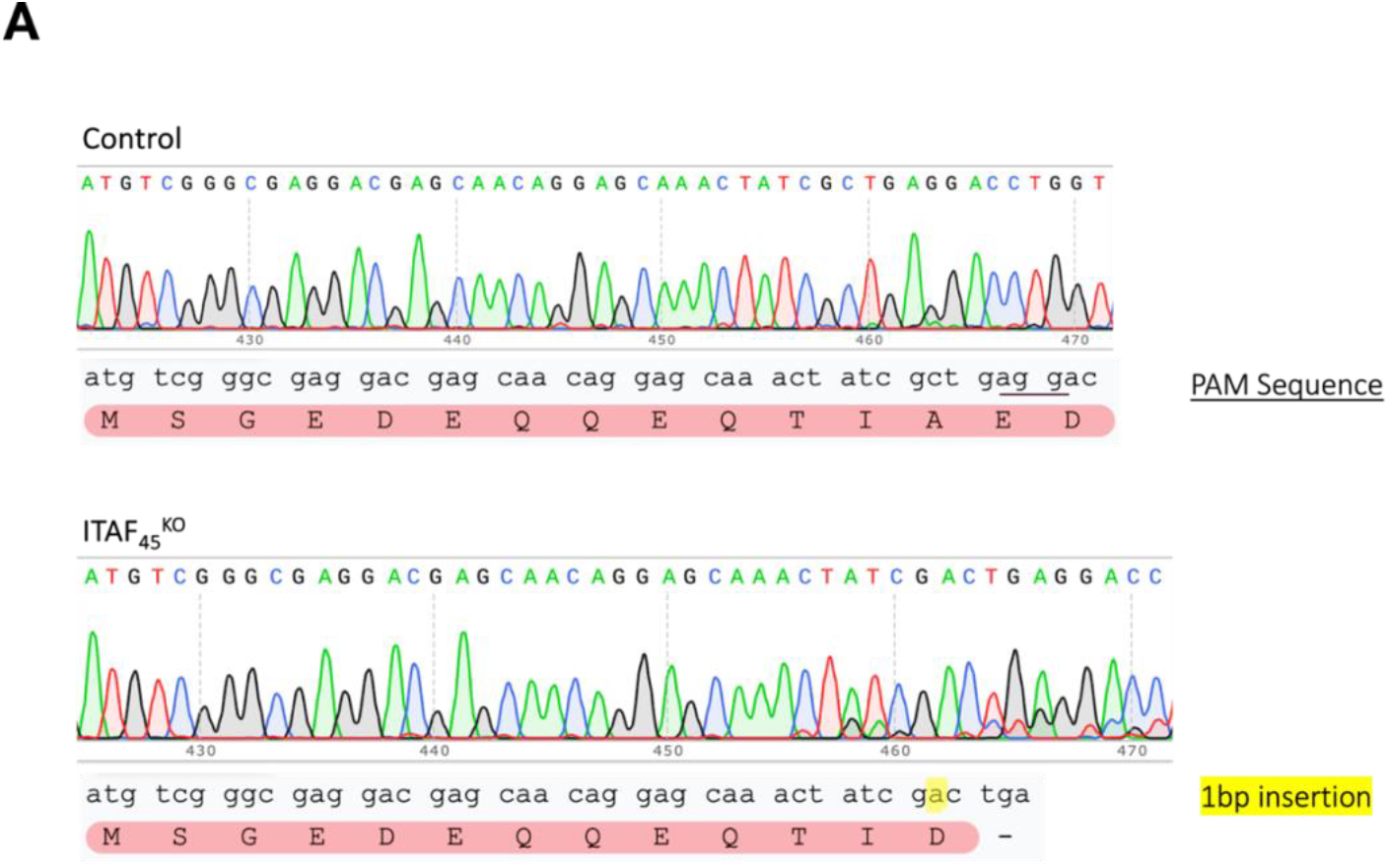
Sanger sequencing validation of ITAF_45_ Knockout. **A**. Chromatogram of H1-HeLa control and ITAF_45_ cells sequences.

**Figure S2.**
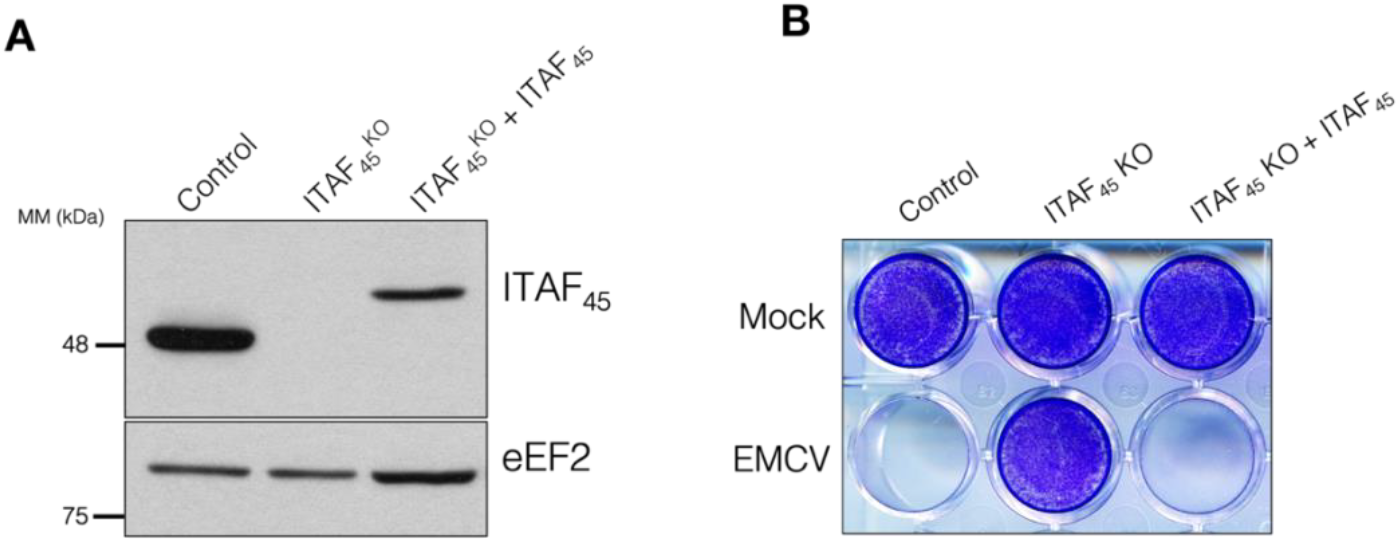
Generation of ITAF_45_ KO in eHAP1 cell lines. **A**. Immunoblot of eHAP1 control, ITAF_45_^KO^, and ITAF_45_^KO^ + MYC-DDK-ITAF_45_ cell lines. **B**. Crystal violet staining of viable cells infected with EMCV (*Mengo* strain) at an MOI of 10 for 24 hrs with “mock” representing uninfected cells.

## Notes

### Competing Interest Statement

The authors have declared no competing interest.

